# Comparative genomic analysis of cyanobacteria as amphibian food sources: insights into high-temperature tolerance potential

**DOI:** 10.1101/2025.08.04.668509

**Authors:** Tetsushi Komoto, Linh Thi Thuy Cao, Haruki Takayanagi, Hinayo Ogino, Haruto Koyama, Satoru Watanabe, Junpei Nomura, Ryuichi Hirota, Takeshi Igawa

## Abstract

Hot-springs represent one of the most accessible yet extreme environments on the Earth’s surface. We isolated and characterized two novel cyanobacterial strains of the genus *Leptolyngbya, L*. sp. Akita and *L*. sp. Seranma from geothermal hot-springs in Japan. These strains showed distinct morphological features; *L*. sp. Akita exhibited linear filamentous structures and blue-green color when cultured under white light, whereas *L*. sp. Seranma exhibited loosely coiled structures and a brown color under the same conditions. Whole-genome sequencing and comprehensive genomic analyses of their genomes identified the acquisition of gene clusters related to metabolic processes, which may be associated with strain-specific phenotypes. In addition, intestinal metagenomic analysis of tadpoles (*Buergeria buergeri* and *B. japonica*) sympatrically living with these strains in the hot-springs suggested that the tadpoles utilized these *Leptolyngbya* as temporal or regular foods in high-temperature environments. These findings provide insights into the ecological significance of these cyanobacteria in extreme environments and their potential applications in biotechnology and the ecological conservation of primary consumers, including amphibians.

## Introduction

Cyanobacteria are essential microorganisms that act as primary producers in aquatic ecosystems and play a pivotal role in sustaining the food web dynamics. In extreme environments, where nutrient and biomass availability is constrained, environmental adaptation and establishment of cyanobacteria are critical events that influence the local ecosystems (Klatt *et al*. 2011; Etesami 2025; Xu *et al*. 2025). Temperature is one of the most fundamental environmental factors to which all organisms, including cyanobacteria, must adapt during evolution and the expansion of their distribution ranges (Paerl and Huisman 2009; Garcia-Pichel *et al*. 2013; Kees *et al*. 2022; Coello-Camba and Agustí 2024).

In the context of climate change, the adaptation and formation of ecosystems under extreme temperature conditions, particularly high temperatures, have garnered increasing attention in recent years. Novel strains of cyanobacteria that have adapted to extremely high-temperature environments have been identified within multiple families, including *Oculatellaceae* (Tang *et al*. 2021), *Leptolyngbyaceae* (Tang *et al*. 2022), *Trichocoleusaceae* (Tang *et al*. 2023), *Thermosynechococcaceae* (Arnold *et al*. 2024), and *Geminocystaceae* (Chen *et al*. 2025). The evolutionary adaptations of cyanobacteria to high-temperature environments not only underscore their resilience but also highlight their potential as bioindicators of the impact of climate change on aquatic ecosystems in the tropics.

As these microorganisms flourish under extreme conditions, they significantly contribute to primary productivity, which is vital for the survival of various aquatic species that depend on them as food. Furthermore, they are increasingly recognized as producers of bioactive nutrient compounds, including proteins, carbohydrates, essential fatty acids, vitamins, minerals, and other functional bioactive compounds, which have potential applications in food or supplements (Vaz *et al*. 2016). Additionally, the metabolic pathways of thermophilic cyanobacteria have evolved to enable the synthesis of unique compounds that can be harnessed for biotechnological applications, thereby broadening their significance beyond their ecological contributions (Rasul *et al*. 2024). As climate change continues to reshape aquatic environments, understanding these adaptations not only informs us about the resilience of cyanobacteria but also emphasizes their potential role in sustainable practices aimed at mitigating the impacts of rising temperatures on ecosystems (Pandey *et al*. 2025).

In a unique ecosystem adapted to high-temperature environments, we recently discovered two homologous cases of ecological symbiosis between cyanobacteria and anuran amphibians of the genus *Buergeria* in natural hot springs in Japan. One of the *Buergeria* species, *B. japonica*, known as the “hot-spring frog,” has tadpoles that thrive at the highest recorded water temperature (46.1°C) in natural hot-springs on Kuchinoshima Island, Kagoshima Prefecture (Komaki, Lau and Igawa 2016). Another species, *B. buergeri*, has also been reported to inhabit a river where a natural hot spring occurs in Kawara-no-yukko, Yuzawa City, Akita Prefecture (Takayanagi 2019). Interestingly, both populations of *Buergeria* species coexist with microbial mats and utilize them as food. Amphibians are crucial vertebrates that serve as omnivores during the tadpole stage and as consumers after metamorphosis within the trophic level. Moreover, nutrient regeneration stemming from both herbivorous and carnivorous feeding activities of tadpoles positively influences the production rates of benthic algae, phytoplankton, and herbivorous benthic chironomid larvae (Iwai, Kagaya and Alford 2012). Consequently, both local ecosystems, initiated by cyanobacteria and expanded by sympatric amphibians, represent intrinsic cases for understanding ecological and evolutionary adaptations and ecosystem formation.

The main goal of this research is to better understand how cyanobacteria adapt to extreme temperatures and the effects of these adaptations on ecosystem stability and dynamics. By examining the symbiotic connections between cyanobacteria and amphibians, especially the genus *Buergeria*, we sought to clarify how these relationships improve nutrient cycling and productivity in harsh environments. In this study, we aimed to clarify the evolutionary genomic adaptation of cyanobacteria and its connection with amphibians living in hot springs. We first isolated two novel cyanobacterial strains from these extreme environments. Subsequently, we determined the whole genome sequences of the isolated strains and conducted comprehensive comparative genomic analyses to identify the specific genes responsible for their remarkable high-temperature tolerance. Additionally, we performed metagenomic analyses of tadpole intestinal microbiota to confirm their dietary reliance on cyanobacterial sources. This multifaceted approach not only enhances our understanding of the ecological dynamics at play but also sheds light on the evolutionary adaptations that enable both cyanobacteria and amphibians to thrive in extreme thermal conditions.

## Materials and methods

### Sample collection, isolation, and growth conditions

Cyanobacteria were isolated from microbial mats collected at two geothermal hot-spring sites in Japan: one from “Kawara-no-yukko” (hot-spring in the Yakunaigawa River), Akinomiya Akita Pref. in August 2022 (hereafter referred to as Akita) and the other from Seranma hot-spring, Kuchinoshima Island, Kagoshima Pref. in June 2023 (hereafter referred to as Seranma). At both sites, microbial mats were found in small, warm water streams originating from the hot-springs and eaten by tadpoles of frogs which belong to the genus *Buergeria* (*B. buergeri* in “Kawara-no-yukko” and *B. japonica* in Seranma hot-spring). We collected the mats using sterile forceps and spoons and immediately transferred them into 50 mL Falcon tubes filled with local hot-spring water. During transport to the laboratory, the samples were kept cool (approximately 10 °C) and in the dark to maintain microbial viability. In the laboratory, the samples were gently rinsed with sterile distilled water to remove loosely attached sediment and non-target debris. Small fragments of the mat were inoculated into 200 mL Erlenmeyer flasks containing 50 mL sterile BG11 liquid medium. These flasks were incubated at 34 °C in a growth chamber (PHCbi, MLR-352, Japan) equipped with 37 W fluorescent lamps (FL40SSENW37HF3, Panasonic, Japan), with the light intensity set to 50 μmol photons/m^2^/s to enrich phototrophic cyanobacteria. After several days of cultivation, cultures displaying characteristic green-brown pigmentation were subcultured two to three times in fresh BG11 medium to reduce microbial diversity and further enrich the cyanobacterial populations. Following enrichment, aliquots of the liquid cultures were spread onto BG11 agar plates and incubated at 40 °C under continuous illumination to obtain individual cyanobacterial colonies. After several days of incubation, filamentous and aggregated colonies appeared on the plates. To eliminate bacterial contamination and obtain axenic cultures, the streaking was repeated multiple times on fresh BG11 agar plates. Colonies exhibiting consistent morphologies were transferred to liquid BG11 medium for further purification.

Complementary chromatic acclimation experiments were performed using a multi-cultivator system (MC-1000; Photon Systems Instruments, Czech Republic) equipped with programmable light-emitting diode (LED) light modules. Cyanobacterial cultures were incubated at 40 °C under continuous illumination of white, red, or green LED light, each set to a photosynthetic photon flux density (PPFD) of 60□μmol photons□m□^2^□s□^1^. The peak emission wavelengths of the red and green LEDs were approximately 615 nm and 530 nm, respectively. Absorption spectra of intact cells were measured using a UV–visible spectrophotometer and normalized to OD680.

### Genomic DNA extraction and sequencing of two cyanobacteria isolates

Genomic DNA from two purified cyanobacterial cultures was extracted for whole-genome sequencing using the DNeasy PowerBiofilm Kit (Qiagen, Hilden, Germany), and libraries for the Oxford Nanopore Technology (ONT) sequencer were constructed using a ligation library preparation kit (SQK-LSK114, ONT). All procedures were performed according to the manufacturer’s instructions, and the quality and molecular weight of genomic DNA were measured using a Qubit fluorometric quantification system (Thermo Fisher Scientific, Inc.). Sequencing was conducted using a MinION sequencer (MinION Mk1B, ONT) and a Flongle flow cell (FLO-FLG114, ONT). The raw sequence data were base-called using a super-accuracy model with duplex base calling, utilizing Dorado (version 0.9.5) and duplex tools (version 0.3.3).

Genome assembly, assessment of completeness, gene prediction, and annotation Sequenced reads with a q-score of > 10 were assembled using Flye (v2.9.4-b1799) (Kolmogorov *et al*. 2019) with the “--nano-hq” parameter. Genome completeness was assessed using the Benchmarking Universal Single-Copy Orthologs (BUSCO (version: 5.7.0)) gene set (cyanobacteria_odb10) (Manni *et al*. 2021). For these two assembled genome sequences and published genomic FASTA files of other *Leptolyngbya* spp. (Table S1) were used for gene prediction and annotation using the prokaryotic gene annotation pipeline PGAP (version: 2024-07-18. build7555) (Tatusova *et al*. 2016). Functional domains in each gene were estimated using InterProScan (version 5.59-91.0) (Jones *et al*. 2014), and Pfam IDs obtained from InterProScan were translated to GO terms using pfam2go in ragp (version 0.3.5.9000) (Dragićević *et al*. 2021). The *rfpA, rfpB*, and *rfpC* genes in *L*. sp. Seranma and *L*. sp. JSC-1 were identified based on PGAP annotations. Genomic regions corresponding to the *rfpABC* cluster and adjacent genes were extracted from the genomes of *L*. sp. Seranma and *L*. sp. JSC-1. To assess gene content, order, and orientation within the *rfpABC* loci, visualization of gene cluster organization and calculation of sequence similarity were conducted using Geneious (version: 9.1.8) (Kearse *et al*. 2012)

### Molecular phylogenetic analyses using 16S ribosomal RNA (rRNA) and Unicore gene set

First, to infer the phylogenetic placement of our isolated cultured strains in the phylum Cyanobacteriota, we analyzed the 16S rRNA sequences of strains Akita and Seranma using Cydrasil (Roush, Giraldo-Silva and Garcia-Pichel 2021). We then conducted genus-level phylogenetic analyses using 16S rRNAs and Unicore (universal and core) genes (single-copy core genes common to most members of a clade) of *Leptolyngbya* spp. In the 16S rRNA analyses, to exclude truncated copies of 16S rRNA, we selected copies 1,400 bp or more in length in each strain. We could not find 16S rRNA sequences of 1,400 bp or more in *L*. sp. JSC-1, a sequence deposited in the NCBI database (accession ID: FJ788926), was used. After multiple alignments of 16S rRNA sequences using MAFFT (version: 7.525, parameters: --auto) (Katoh and Standley 2013), a phylogenetic tree was constructed using IQ-TREE (version: 2.4.0, parameters: -m MFP - B 1000) (Minh *et al*. 2020). In the analyses of the Unicore gene set, single-copy genes in more than 80% of all strains of *Leptolyngbya* spp. were extracted. Using the pipeline included in the Unicore package (Kim, Park and Steinegger 2024), this gene set was concatenated, aligned, and used for phylogenetic tree construction. These phylogenetic trees with bootstrap values were visualized as unrooted trees using iTOL (https://itol.embl.de/) (Letunic and Bork 2024).

Synteny identification and comparative genome analysis using predicted gene models We constructed synteny plots between *L*. sp. Akita and *L*. sp. Seranma, *L*. sp., and their phylogenetically closest strains using ReCCO (version: 1.0-beta, download link of script; https://www.genome.jp/ftp/tools/ReCCO/ReCCO_Readme.html), and the plots were visualized using DiGAlign (version: 2.0, web server; https://www.genome.jp/digalign/) (Nishimura *et al*. 2024) with default parameters and small modifications for proper execution.

For comparative genomic analyses, reciprocal BLASTP (version 2.15.0) (Altschup *et al*. 1990) searches were performed between the predicted genes of strain Akita, strain Seranma, and *L*. sp. JSC-1 and predicted genes of each strain of *Leptolyngbya* spp.. We classified the genes of strains Akita, Seranma, and *L*. sp. JSC-1 into three categories: the gene was a reciprocal best hit (RBH), not the reciprocal but best hit (BH), and no homologous genes (NH). We used e-value ≤ 1.0e-10 in these BLASTP searches. After classifying genes into these three categories, hierarchical clustering based on Gower’s distance and Ward’s method was performed using state matrices of strain Akita, strain Seranma, and *L*. sp. JSC-1 was formed by classifying category vectors corresponding to other *Leptolyngbya* strains. To obtain significantly enriched Gene Ontology (GO) terms in each gene cluster, genes were divided into clusters using appropriate thresholds from the results of hierarchical clustering, and Fisher’s exact probability tests with multiplicity adjustment using the Benjamini-Hochberg method were performed for each gene cluster.

### Identification of evolutionally novel heat tolerance-related genes

We compared gene repertoires between hot-spring-derived strains and those from non-thermal environments among evolutionary close strains on the Unicore phylogenetic tree (Fig. 3) to identify genes associated with heat tolerance that were evolutionarily acquired. Based on the topology of the Unicore phylogenetic tree, we searched the genes being RBH within the hot-spring-derived strains (Akita, Seranma, and JSC-1), and NH in the two strains which share the most recent common ancestors with the hot-spring-derived strains in the tree (*L. boryana, L*. sp. GGD, *L. ohadii*, and *L*. sp. FACHB-711) (Fig. 3)

### Metagenomic analysis of tadpole intestinal microbiota using 16S rRNA

To confirm the ecological significance of cyanobacteria in extreme environments, we verified tadpole feeding by metagenomic analysis of tadpole intestinal microbiota using 16S rRNA. We obtained tadpoles of the Japanese bell-ring frog, *B. buergeri* from both “Kawara-no-yukko” in August 2022 and tadpoles of the Ryukyu bell-ring frog, *B. japonica* from Seranma hot-spring in May 2022. Three *B. buergeri* tadpoles from different water temperatures of 26 °C, 28 °C, and 37 °C, and nine *B. japonica* tadpoles from the same water temperature of approximately 40 °C were collected and preserved in 70% ethanol after deep anesthetization using MS-222 and kept in a -30 °C freezer until DNA extraction. All sample collections were approved by the Hiroshima University Animal Research Committee (approval number: G14-2). Total DNA, including intestinal microbial metagenomic DNA and tadpole DNA, was extracted from the tadpole intestine using DNAs-ici.-F (Rizo Inc., Tsukuba, Japan) according to the manufacturer’s instructions and dissolved in 50 μL of nuclease-free water. We amplified full length 16S rRNA gene using 27F (5’-AGAGTTTGATCCTGGCTCAG-3’) and 1492R (5’-TACGGYTACCTTGTTACGACTT-3’). Polymerase chain reaction (PCR) amplification was conducted in a 25 μL reaction with 0.5 U KOD FX Neo (Toyobo Co., Ltd. Osaka, Japan), 1 × PCR Buffer for KOD FX Neo, 0.4 mM each dNTP, 0.2 μM each primer, and 1 μL total DNA samples with the following cycle condition: initial denaturation at 94 °C for 3 min, then 35 cycles of 10 s denaturation at 98 °C, 30 s annealing at 55 °C, and 2 min elongation at 68 °C. The PCR products were then removed from the remaining primers by polyethylene glycol precipitation and dissolved in 5 μL of nuclease-free water. After measuring the concentration using a Qubit Fluorometer (Thermo Fisher Scientific) and Qubit dsDNA HS Assay Kit (Thermo Fisher Scientific), each precipitated sample was used for barcoded library construction for Nanopore sequencing following the native barcoding protocol (SQK-LSK109 with EXP-NBD104 and EXP-NBD114, NBE_9065_v109_revAP_14Aug2019). Following the manufacturer’s instructions, sequencing was conducted using a MinION sequencer (MinION Mk1B, ONT) and Flongle flow cell (FLO-FLG001, ONT). MinKNOW (version: 22.03.6) was used for basecalling and demultiplexing with Super-accuate basecalling model, and the resultant FASTQ files of each sample were and analyzed EPI2MElabs platform (version: 5.2.3) with wf-metagenomics nextflow workflow (version: 2.13.0) and pre-built “core-nt” Kraken2/Bracken database (downloaded 4/27/2025, https://benlangmead.github.io/aws-indexes/k2). Differential abundance analysis was conducted using ALDEx2 (version 1.36.0) (Fernandes *et al*. 2013) packages, which outputs multiple statistical tests, including Welch’s t-test, Wilcoxon rank-sum test, Kruskal-Wallis test, and generalized linear model (GLM) with significance assessed by Wald test, based on centered log-ratio-transformed abundances. The p-values were adjusted using the Benjamini-Hochberg method.

## Results

### Morphological evaluation and whole-genome sequencing of the isolated cyanobacterium strains

After repeated subcultures of microbial mats collected from hot-springs in Kawara-no-yukko and Seranma, fine green and brown cyanobacterial strains were obtained. Both strains exhibited filamentous and non-heterocystous structures under a light microscope (Fig. 1A and 1B). Considering their hot-spring inhabitation and molecular phylogenetic relationships (see next section), we considered both isolated strains as members of the genus *Leptolyngbya* and designated them as *Leptolygbya* sp. Akita and *L*. sp. Seranma, named after the respective collection sites. *L*. sp. Akita showed a relatively linear filamentous structure, whereas *L*. sp. Seranma has a loosely coiled filamentous structure. We obtained 405,522,732 bp and 933,140,049 bp of raw sequence data for *L*. sp. Akita and *L*. sp. Seranma (Table 1). The resultant genome assemblies based on these data consisted of three circular and one linear contig for Akita (6,829,225 bp in total) and two circular and two linear contigs for Seranma (8,049,769 bp in total) (Table 1). The BUSCO completeness values of these genomes were 95.5% and 99.6%, while that only 1.4% and 0.3 % of the missing BUSCO genes, respectively.

**Table 1.** Metrics of the draft genomes of strains Akita and Seranma with BUSCO completeness. Linear and circular contigs are indicated as LC and CC, respectively.

| Strain | Akita | Seranma |
| --- | --- | --- |
| Total number of reads | 98,144 | 163,820 |
| Total length of reads (bp) | 405,522,732 | 933,140,049 |
| N50 of reads (bp) | 17,945 | 6,502 |
| > Q20 (%) | 65.18 | 81.28 |
| > Q30 (%) | 35.04 | 67.18 |
| Total number of contigs | 4 | 4 |
| Total assembly size (bp) | 6,829,225 | 8,049,769 |
| GC content (%) | 46.75 | 50.77 |
| Assembly size and state of contigs (bp) | contig_1: LC, 277,981<br>contig_3: CC, 6,300,885<br>contig_4: CC, 10,498<br>contig_5: CC, 239,861 | contig_1: LC, 1,313,761<br>contig_2: CC, 6,476,496<br>contig_3: LC, 226,323<br>contig_4: CC, 33,189 |
| BUSCO completeness (% , cyanobacteria_odb10, 773 genes from 141 genomes) | Complete: 95.5<br>(Single-copy: 95.2, Duplicated: 0.3)<br>Fragmented: 3.1<br>Missing: 1.4 | Complete: 99.6<br>(Single-copy: 98.6, Duplicated: 1.0)<br>Fragmented: 0.1<br>Missing: 0.3 |
| Total number of estimated genes from PGAP (genes) | 6,307 | 6,752 |

**Fig. 1.**
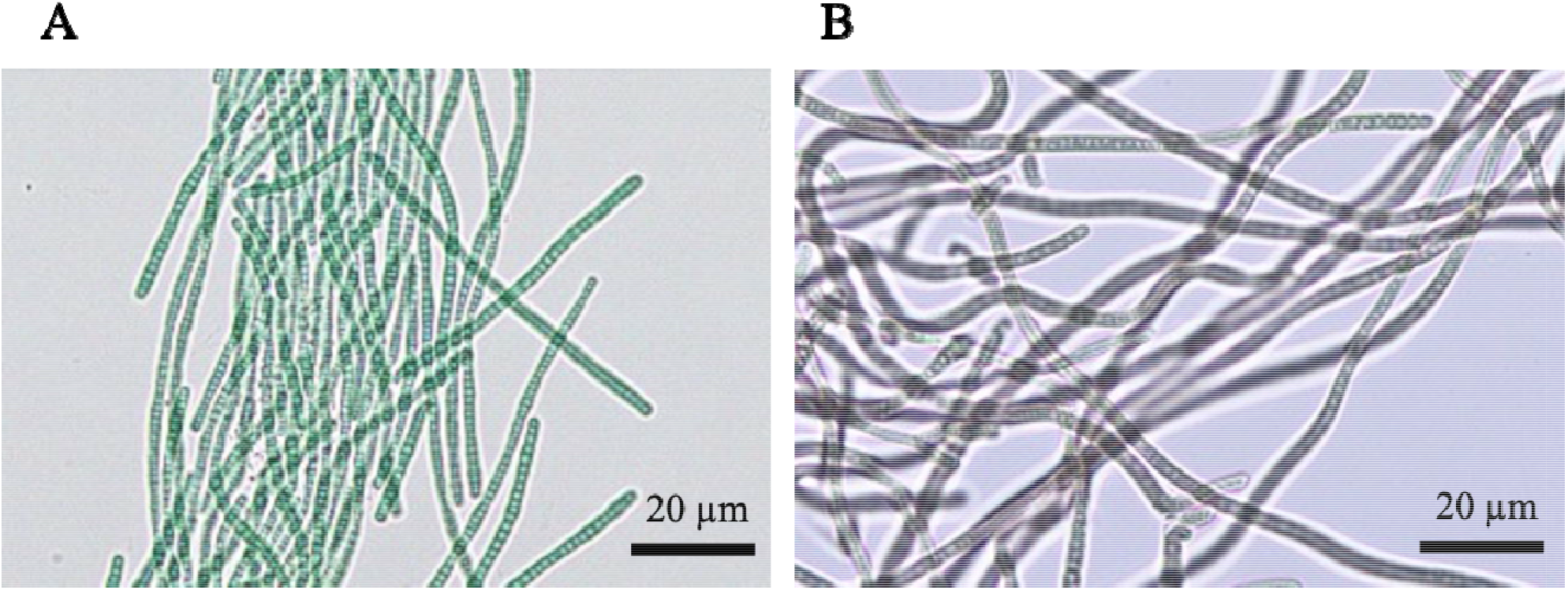
Photographs of the novel isolated strains. Microphotographs of (A) *L*. sp. Akita, and (B) *L*. sp. Seranma.

Light-dependent absorption spectral changes and conservation of the *rfpABC* gene cluster Absorption spectra of cells grown under different light conditions were measured for *L*. sp. Akita and *L*. sp. Seranma following acclimation to monochromatic light at 615 nm (red light) or 530 nm (green light), as well as to white light conditions. *L*. sp. Seranma exhibits complementary chromatic acclimation, appearing brown under white light and green under red light (Fig. 2A), similar to *L*. sp. JSC-1 (Gan *et al*. 2014).

**Fig. 2.**
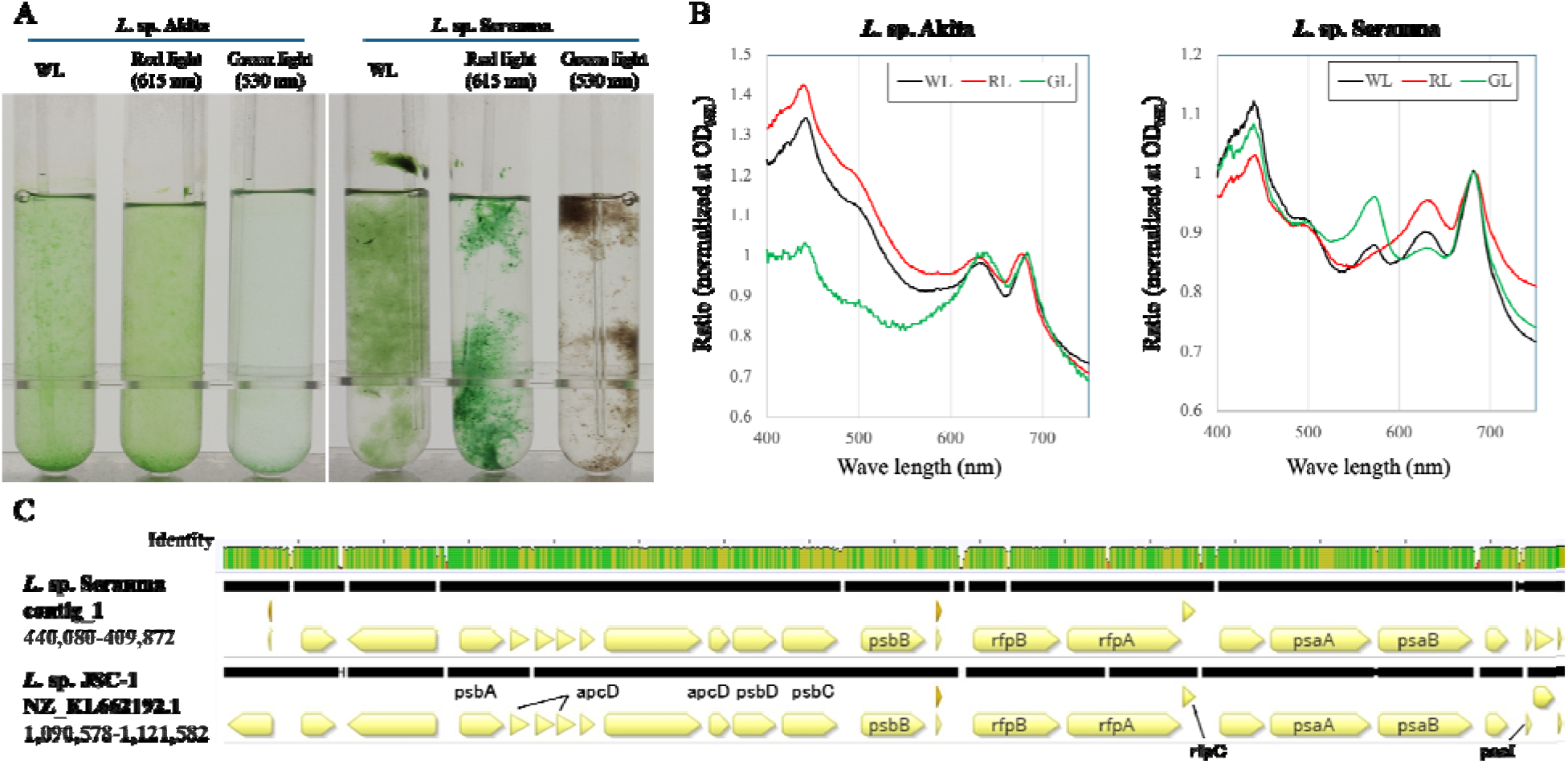
Coloration of *L*. sp. Akita and *L*. sp. Seranma grown under white, red, and green light conditions (A), corresponding absorption spectra (B). Comparative genomic structure and sequence similarity of the *rfp* gene cluster and surrounding regions (C). Sequence alignment of the flanking regions was visualized using Geneious, with identity values displayed according to Geneious.

In *L*. sp. Akita, the distribution of the absorption spectra remained largely unchanged across the different light conditions (Fig. 2B). In particular, the positions of the major absorption peaks showed minimal variation. In contrast, *L*. sp. Seranma exhibited pronounced light-dependent changes in its absorption spectra. Cells acclimated to different light conditions displayed distinct shifts in the spectral distribution, resulting in absorption profiles (Fig. 2B) that closely resembled those reported for far-red light– acclimated cyanobacteria by Gan *et al*.

In addition, previous studies have demonstrated that *rfpA, rfpB*, and *rfpC* function as master regulatory elements controlling far-red light photoacclimation (FaRLiP) and are encoded within a conserved gene cluster (Fig. 1A in Gan et al., 2014). To assess whether the differential spectral responses observed between *L*. sp. Seranma and *L*. sp. JSC-1 could be associated with the presence and genomic organization of *rfp* genes, we examined the genomic regions surrounding *rfpA, rfpB*, and *rfpC* in both strains. In *L*. sp. JSC-1, a genomic region encompassing *rfpA, rfpB*, and *rfpC* (pgaptmp_006202, pgaptmp_006201, and pgaptmp_006203, respectively) was extracted from NZ_KL662192.1 (positions 1,090,578–1,121,582). This region was compared with the corresponding *rfp* locus in *L*. sp. Seranma (pgaptmp_000321, pgaptmp_000322, and pgaptmp_000320) located on contig_1 (positions 440,080-409,872). Comparative analysis revealed that the genomic organization of the *rfpABC* cluster was highly conserved between *L*. sp. Seranma and *L*. sp. JSC-1. Not only were the three *rfp* genes preserved in the same order and orientation, but the surrounding genes also exhibited a conserved synteny across the two strains.

### Phylogenetic relationships in cyanobacteria and genus *Leptolyngbya*

The phylogenetic tree of 16S rRNA generated by Cydrasil showed that both isolated cultured strains were in a branch containing most of the *Leptolyngbya* sp. (Fig. S1). We then inferred the evolutionary relationships of the two strains and known strains of *Leptolyngbya* (except for the strains with extremely low BUSCO completeness) using 16S rRNA and the Unicore dataset of *Leptolyngbya* spp. Based on the PGAP annotation, 75 copies of 16S rRNA were identified in the *Leptolyngbya* spp. genomes, and 48 of these (including a manually added sequence of *L*. sp. JSC-1) were extracted as valid 16S rRNA and used for phylogenetic analysis. In the phylogenetic tree of 16S rRNA, *L*. sp. Akita was placed in a cluster containing *L. boryana, L*. sp. FACHB-161, *L*. sp. FACHB-238, *L*. sp. FACHB-239, *L*. sp. FACHB-402, and *L*. sp. GGD, whereas all copies of *L*. sp. Seranma were placed in clusters containing *L*. sp. JSC-1 (Fig. 3A). However, the evolutionary relationships between the strains within the clusters were unclear.

**Fig. 3.**
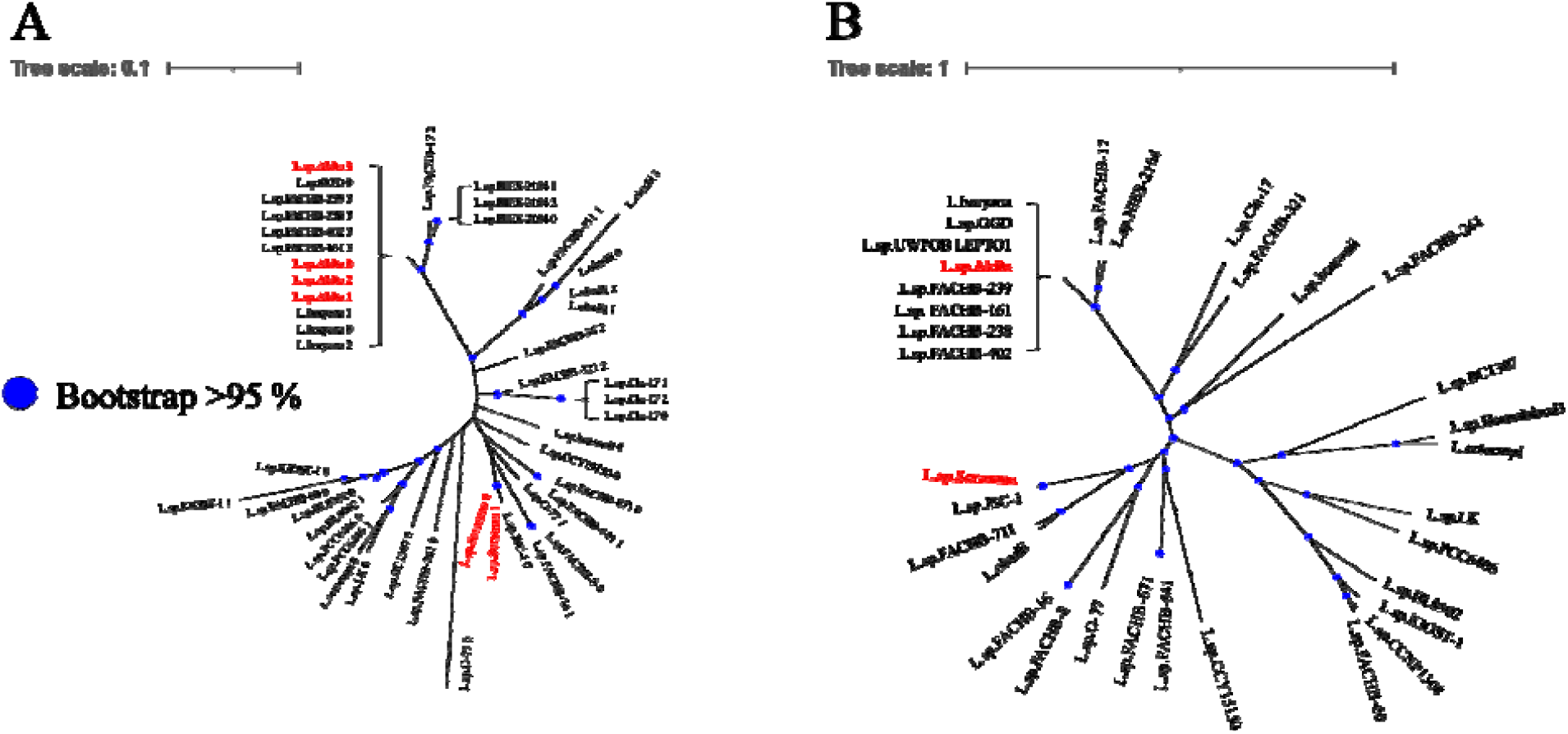
Unrooted phylogenetic trees highlighting internal nodes with bootstrap values >95 % in *Leptolyngbya* spp.. (A) 16S rRNA and (B) Unicore gene set

Using the Unicore pipeline, 408 proteins were selected as the Unicore gene set, and multiple sequence alignment resulted in 100,760 amino acids (including gaps). The phylogenetic tree of the Unicore gene set showed a clustering pattern similar to that of 16S rRNA and clear evolutionary relationships among the strains within the clusters. In the Unicore tree, *L*. sp. GGD was the strain closest to *L*. sp. Akita and *L*. sp. JSC-1 was the strain closest to *L*. sp. Seranma (Fig. 3B).

### Synteny conservation patterns between *Leptolyngbya* spp

We conducted synteny analyses in combination with the closest strains, *L*. sp. Akita - *L. boryana* and *L*. sp. Seranma - *L*. sp. JSC-1. The resultant synteny plots showed macro synteny contig_2 of *L*. sp. Akita and the chromosome of *L. boryana* (NZ_CP130144.1) with 95–100% identity and two large genomic inversions of approximately 0.7–2 Mb (Fig. 4A). Interestingly, *L. boryana* had few homologous genomic regions with contig_1 (approximately 0.3 Mb, 5’ end in Fig. 4A) in *L*. sp. Akita. *L*. sp. Seranma and *L*. sp. JSC-1 also showed macro-synteny over the entire length of contig_2 in *L*. sp. Seranma and the scaffold of *L*. sp. JSC-1 (NZ_KL662191.1) with an identity of 95–100%, except for three genomic inversions of approximately 0.4 Mb and four genomic transpositions of approximately 40–80 kb (Fig. 4B). In addition, the other scaffold in *L*. sp. JSC-1 (NZ_KL662192.1) showed high homology with contig_1 of *L*. sp. Seranma.

**Fig. 4.**
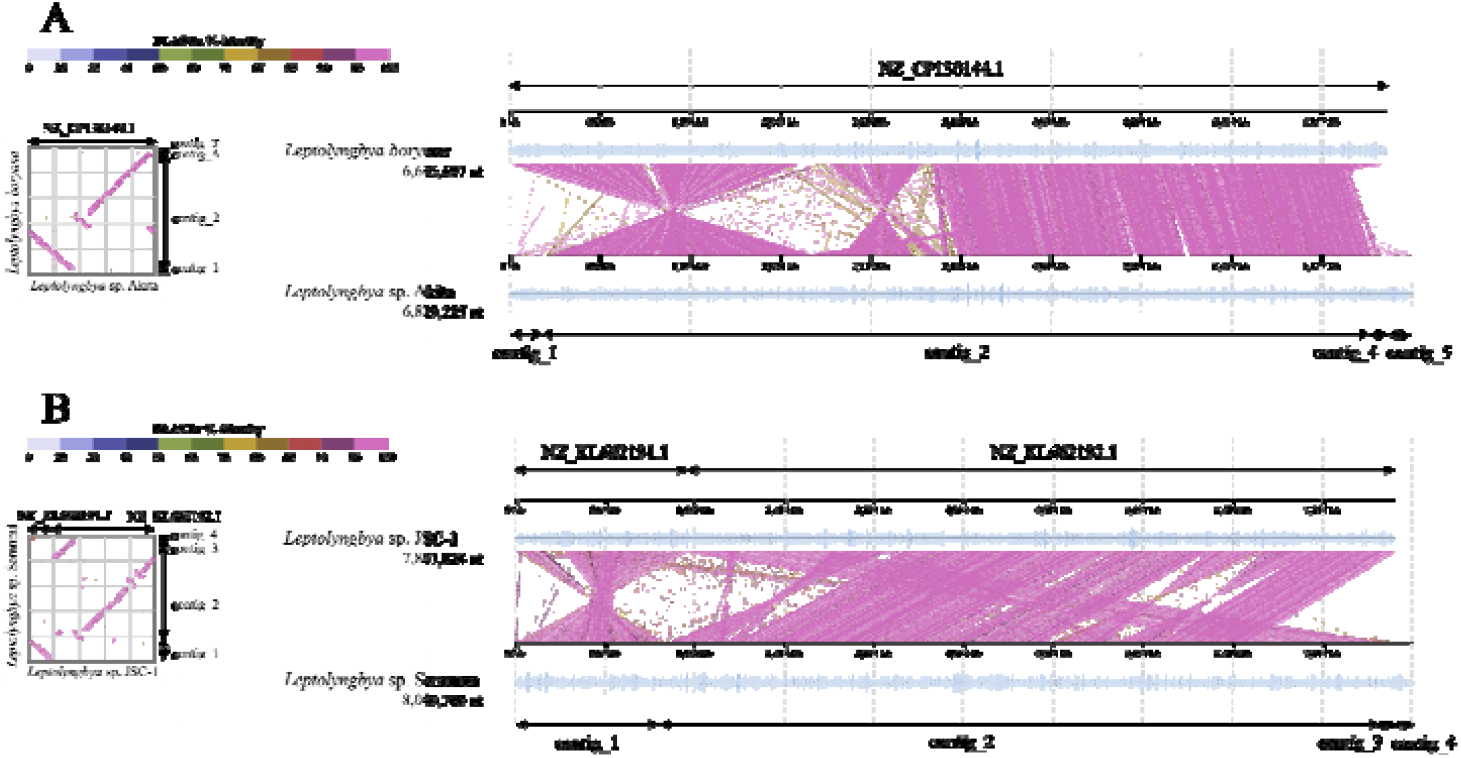
Synteny and dot plots between pairs of *Leptolyngbya* strains. (A) *L*. sp. Akita and *L. boryana* (B) *L*. sp. Seranma and *L*. sp. JSC-1

**Fig. 5.**
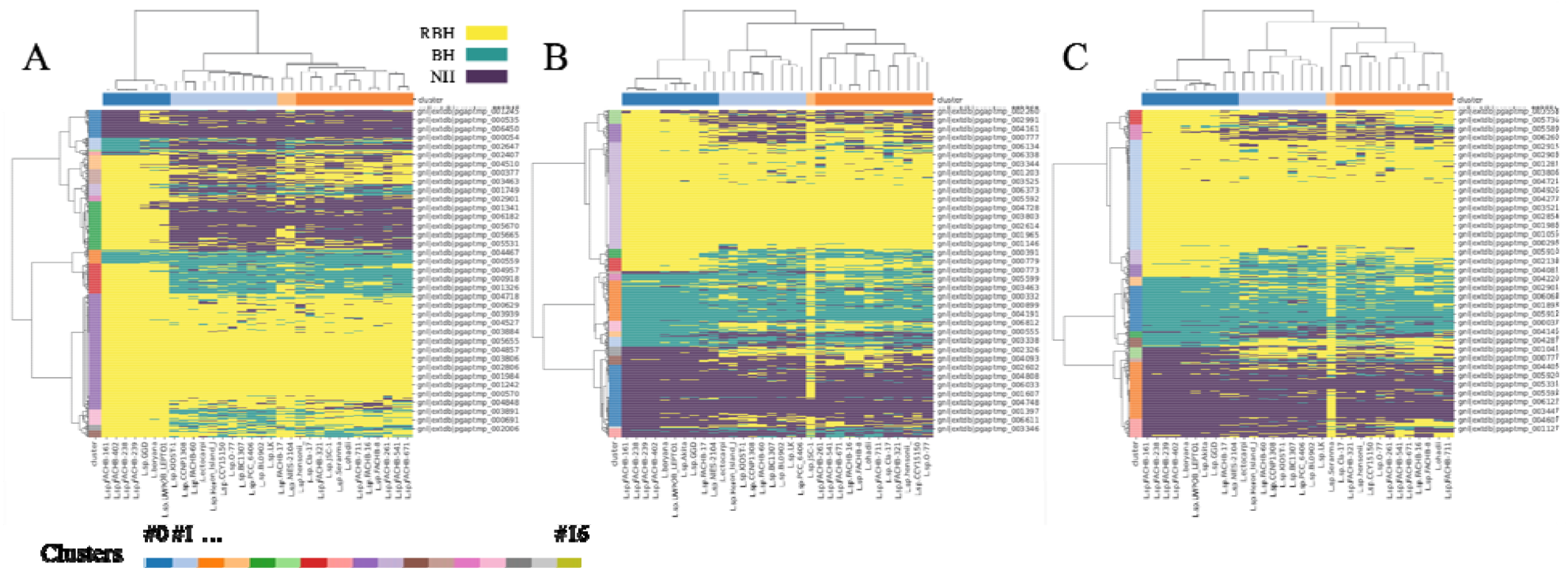
Matrices showing reciprocal BLASTP search between genes of the target strain and genes of the other *Leptolyngbya* strains with dendrograms. The target strains were (A) *L*. sp. Akita; (B) *L*. sp. Seranma, and (C) *L*. sp. JSC-1, respectively. Genes or strains were sorted and colored according to their clusters. See Tables S9-S11 for enriched GO terms in each cluster.

### Strain- or strain-group-specific gene enrichment and functional estimation based on comparative genomic analyses

Based on the PGAP annotation pipeline, 6,307, 6,752, and 6,354 genes were predicted in the genome sequences of *L*. sp. Akita, *L*. sp. Seranma, and *L*. sp. JSC-1 (see Table S1 for predicted genes, including other *Leptolyngbya* strains). Reciprocal BLASTP searches were performed between the predicted genes of *L*. sp. Akita (Table S2) (or *L*. sp. Seranma (Table S3) or *L*. sp. JSC-1 (Table S4)) and the predicted genes of each strain of *Leptolyngbya* spp.. Hierarchical clustering was performed using the BLASTP hit category matrices of *L*. sp. Akita, *L*. sp. Seranma, and *L*. sp. JSC-1. From the dendrograms obtained by hierarchical clustering, all predicted genes of *L*. sp. Akita, *L*. sp. Seranma, and *L*. sp. JSC-1 strains were divided into 15, 13, and 12 gene clusters by selecting appropriate threshold values (Table S5, Fig. S2). Enrichment analyses were performed for the strain-specific gene clusters, which contained genes that specifically accumulated in the target strains and found few homologs in the other strains. These clusters necessarily accumulated the highest number of NH genes, and GO terms, “nitrogen compound metabolic process” and “transition metal ion binding” were identified as significantly enriched terms (q-value (false discovery rate) <0.05 and odds ratio >1) in gene cluster #0 of *L*. sp. Akita (Tables 2A and S9). In addition, “DNA transposition” and “transposase activity” were identified as significantly enriched terms in gene cluster #0 of *L*. sp. Seranma (Table 2B, Table S10). However, no significantly enriched GO terms were identified in gene cluster #2 of *L*. sp. JSC-1 (Table S11). Instead, “phycobilisome” and “photosynthesis”-related GO terms were identified as significantly enriched in gene cluster #2 of *L*. sp. Seranma (Table 2C, Table S10) and gene cluster #0 of *L*. sp. JSC-1 (Table 2D, Table S11). The genes included in these clusters tended to be orthologs because they were only found in synteny blocks of these closely related strains (*L*. sp. Seranma and *L*. sp. JSC-1), but may not be present in other strains.

**Table 2.**
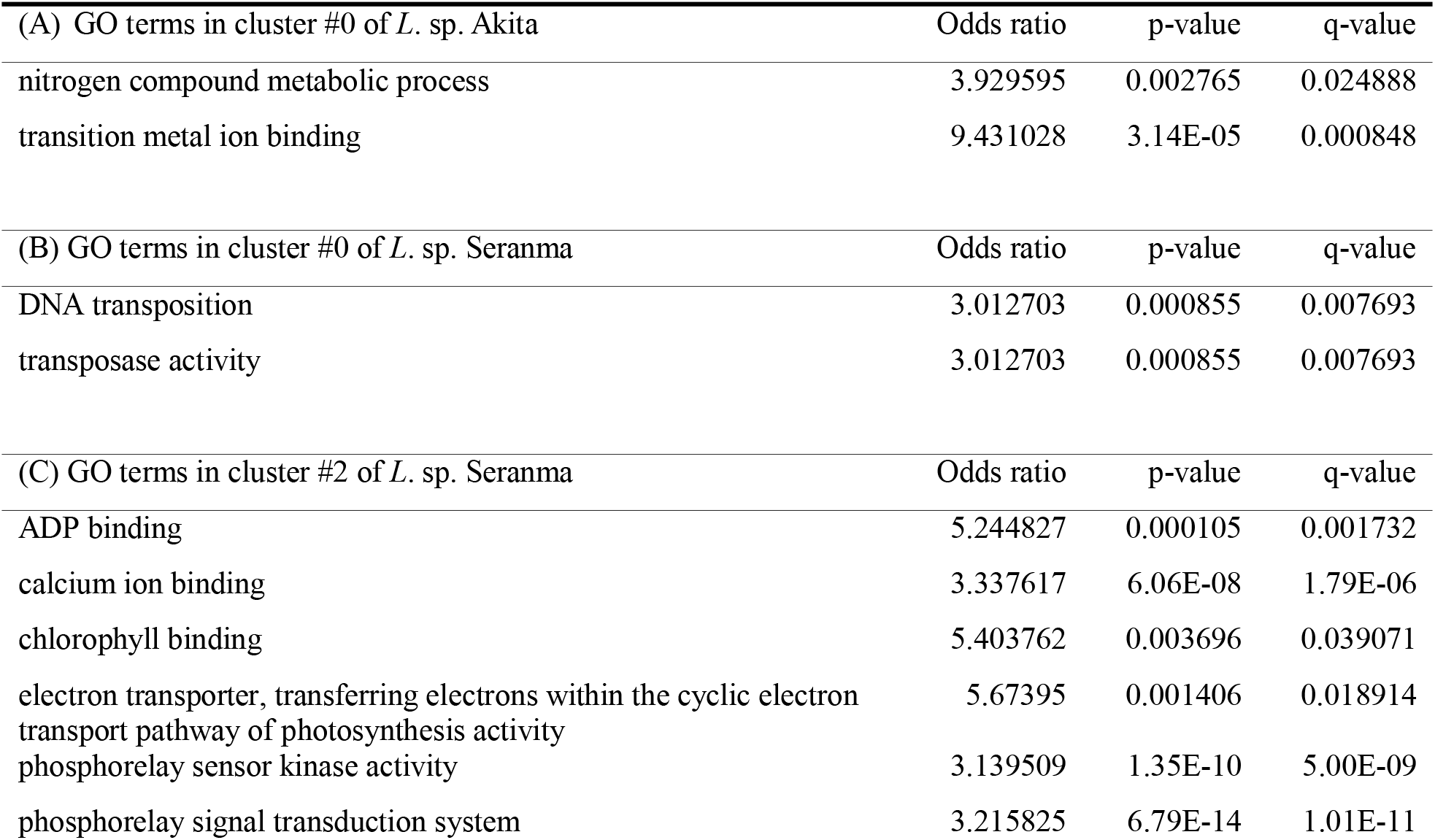
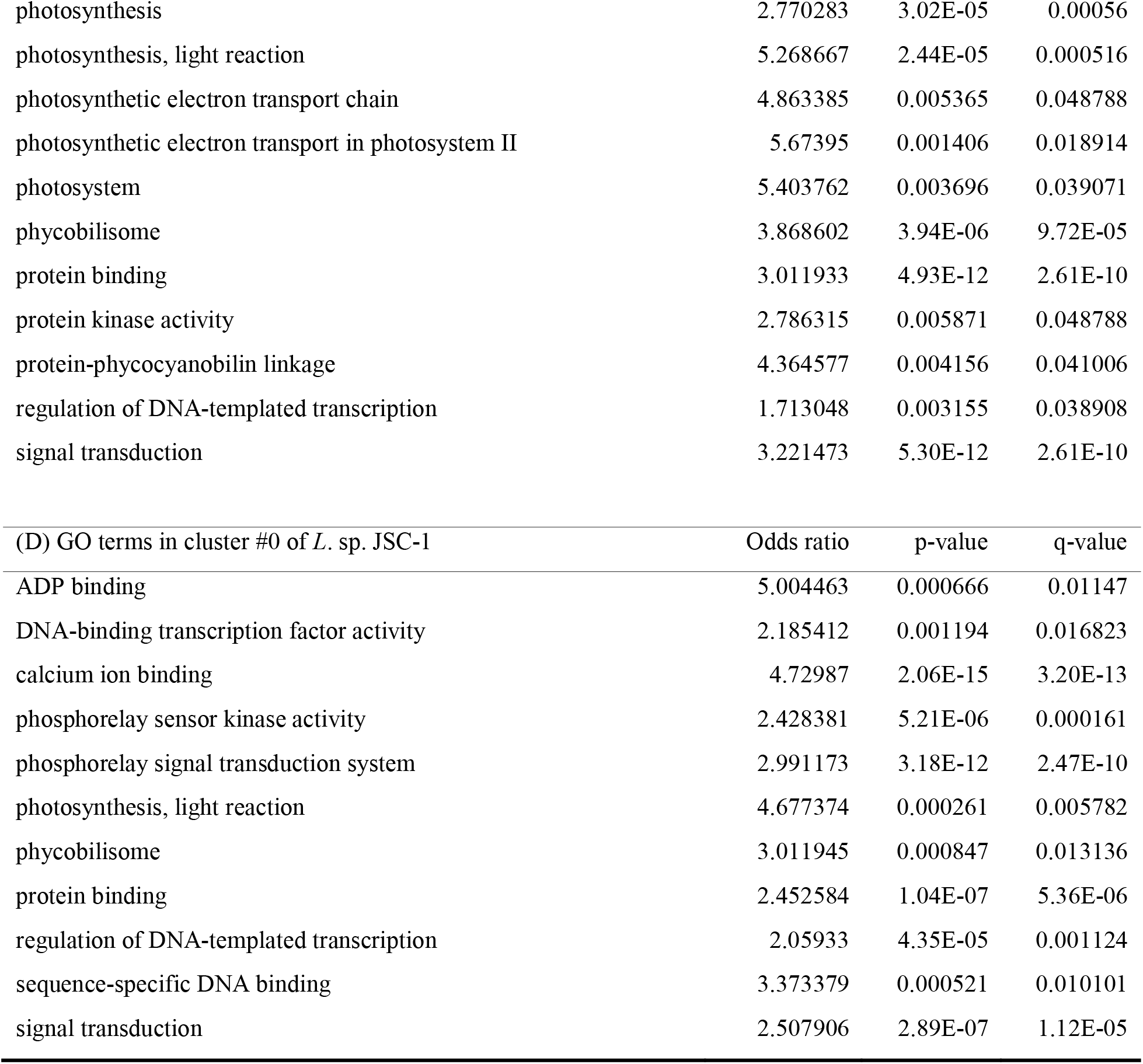
Enriched statistically significant GO terms.

| (A) GO terms in cluster #0 of *L. sp. Akita* | Odds ratio | p-value | q-value |
| --- | --- | --- | --- |
| nitrogen compound metabolic process | 3.929595 | 0.002765 | 0.024888 |
| transition metal ion binding | 9.431028 | 3.14E-05 | 0.000848 |
| (B) GO terms in cluster #0 of *L. sp. Seranma* | Odds ratio | p-value | q-value |
| DNA transposition | 3.012703 | 0.000855 | 0.007693 |
| transposase activity | 3.012703 | 0.000855 | 0.007693 |
| (C) GO terms in cluster #2 of *L. sp. Seranma* | Odds ratio | p-value | q-value |
| ADP binding | 5.244827 | 0.000105 | 0.001732 |
| calcium ion binding | 3.337617 | 6.06E-08 | 1.79E-06 |
| chlorophyll binding | 5.403762 | 0.003696 | 0.039071 |
| electron transporter, transferring electrons within the cyclic electron transport pathway of photosynthesis activity | 5.67395 | 0.001406 | 0.018914 |
| phosphorelay sensor kinase activity | 3.139509 | 1.35E-10 | 5.00E-09 |
| phosphorelay signal transduction system | 3.215825 | 6.79E-14 | 1.01E-11 |
| photosynthesis | 2.770283 | 3.02E-05 | 0.00056 |
| photosynthesis, light reaction | 5.268667 | 2.44E-05 | 0.000516 |
| photosynthetic electron transport chain | 4.863385 | 0.005365 | 0.048788 |
| photosynthetic electron transport in photosystem II | 5.67395 | 0.001406 | 0.018914 |
| photosystem | 5.403762 | 0.003696 | 0.039071 |
| phycobilisome | 3.868602 | 3.94E-06 | 9.72E-05 |
| protein binding | 3.011933 | 4.93E-12 | 2.61E-10 |
| protein kinase activity | 2.786315 | 0.005871 | 0.048788 |
| protein-phycocyanobilin linkage | 4.364577 | 0.004156 | 0.041006 |
| regulation of DNA-templated transcription | 1.713048 | 0.003155 | 0.038908 |
| signal transduction | 3.221473 | 5.30E-12 | 2.61E-10 |

Table 2. Enriched statistically significant GO terms.
| (D) GO terms in cluster #0 of *L. sp.* JSC-1 | Odds ratio | p-value | q-value |
| --- | --- | --- | --- |
| ADP binding | 5.004463 | 0.000666 | 0.01147 |
| DNA-binding transcription factor activity | 2.185412 | 0.001194 | 0.016823 |
| calcium ion binding | 4.72987 | 2.06E-15 | 3.20E-13 |
| phosphorelay sensor kinase activity | 2.428381 | 5.21E-06 | 0.000161 |
| phosphorelay signal transduction system | 2.991173 | 3.18E-12 | 2.47E-10 |
| photosynthesis, light reaction | 4.677374 | 0.000261 | 0.005782 |
| phycobilisome | 3.011945 | 0.000847 | 0.013136 |
| protein binding | 2.452584 | 1.04E-07 | 5.36E-06 |
| regulation of DNA-templated transcription | 2.05933 | 4.35E-05 | 0.001124 |
| sequence-specific DNA binding | 3.373379 | 0.000521 | 0.010101 |
| signal transduction | 2.507906 | 2.89E-07 | 1.12E-05 |

### Putative genes associated with thermal stress adaptation newly acquired by *L*. sp. sp. Akita and the *L*. sp. Seranma–*L*. sp. JSC-1 lineage

Among the PGAP annotations, those mapped to genes that were RBH in hot-spring-derived strains and NH in non-thermal environment-derived strains comprised 117 annotations in *L*. sp. Akita and 386 annotations in *L*. sp. Seranma, respectively. Of these, 27 annotations were detected in both strains, including those potentially related to heat stress response, such as oxidative stress response factors (e.g., peroxiredoxin family protein and Dyp-type peroxidase) and membrane stress response–related factors (e.g., M23 family metallopeptidase, NfeD family protein, and dynamin family protein). In addition to these oxidative and membrane stress response factors, RNA remodeling factors, such as DEAD/DEAH box helicases, were also included (Table 3A).

**Table 3.**
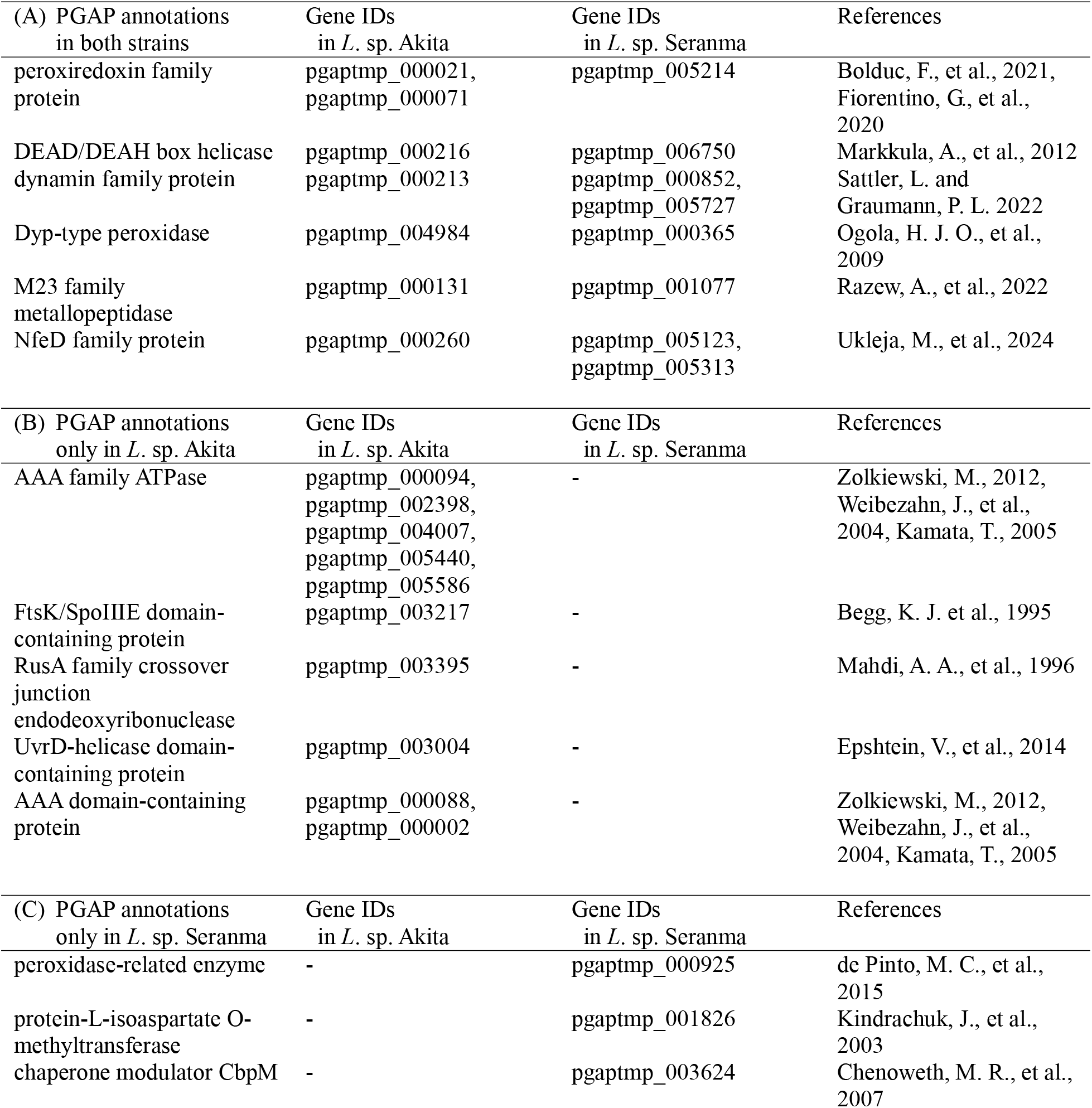
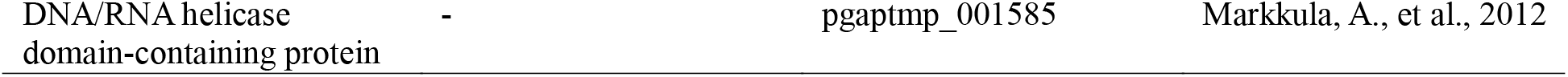
Summary of annotations potentially associated with thermal tolerance and their corresponding reference literature. Annotations not listed in this table are provided in Tables S12-S14.

Furthermore, 90 and 296 annotations were detected exclusively in *L*. sp. Akita and *L*. sp. Seranma, respectively. Annotations were detected only in *L*. sp. Akita included factors involved in DNA protection mechanisms, such as FtsK/SpoIIIE domain-containing proteins, RusA family crossover junction endodeoxyribonucleases, and UvrD helicase domain-containing proteins, as well as protein quality control–related factors involved in disaggregation and proteolysis, such as AAA family ATPases and AAA domain-containing proteins (Table 3B). In contrast, annotations were detected only in *L*. sp. Seranma included oxidative stress response factors, such as peroxidase-related enzymes, protein repair enzymes, such as protein-L-isoaspartate O-methyltransferase, chaperone regulatory factors, such as chaperone modulator CbpM, and DNA/RNA remodeling factors, such as DNA/RNA helicase domain-containing proteins (Table 3C).

Comparison of intestinal microbiota of *B. buergeri* and *B. japonica* tadpoles Intestinal metagenomic analysis of *B. buergeri* and *B. japonica* tadpoles detected 351 genera with non-zero reads using the Kraken2/Bracken pipeline. *Leptolyngbya* was detected in the intestine of *B. buergeri* at a maximum of 1.3%, and was not detected in all tadpoles derived from 37°C. In contrast, *Leptolyngbya* was detected in all intestines of *B. japonica*, accounting for a maximum of approximately 21% (Fig. 6). We then performed a statistical analysis using ALDEx2 to test which genera showed significant differences in variances or abundances among tadpole groups. The results of the Kruskal-Wallis tests showed that there was no significant abundance (q-value >0.05) of any genera between *B. buergeri* tadpoles derived from different temperatures (26, 28, and 37 °C) and those of *B. japonica* tadpoles derived from 40 °C. In contrast, the results of the GLM tests showed statistically significant variance (q-value <0.05) for *Leptolyngbya* and *Limnothrix* (Table S15). The results of Welch’s t-test showed that there was no significant difference in abundance in any comparative pairs of tadpole groups (“all *B. buergeri* tadpoles and all *B. japonica* tadpoles”, “*B. buergeri* tadpoles derived from 26 °C and *B. buergeri* tadpoles derived from 28 °C”, “*B. buergeri* tadpoles derived from 26 °C and *B. buergeri* tadpoles derived from 37 °C”, and “*B. buergeri* tadpoles derived from 28 °C and *B. buergeri* tadpoles derived from 37 °C”). In contrast, the results of Wilcoxon tests on these pairs of tadpole groups showed significant abundance of *Azospira, Cephaleuros, Citrobacter, Enterobacter, Escherichia, Leptolyngbya, Mycobacterium, Oryzias, Pseudophryne, Rhizobium, Streptococcus, Tribolium*, and *Trichoderma* (Tables S16-S19), except for the vertebrate genera possibly derived from the host or contamination. *Leptolyngbya* was detected as a unique significantly abundant genus only in the pair of “all *B. buergeri* tadpoles and all *B. japonica* tadpoles”.

**Fig. 6.**
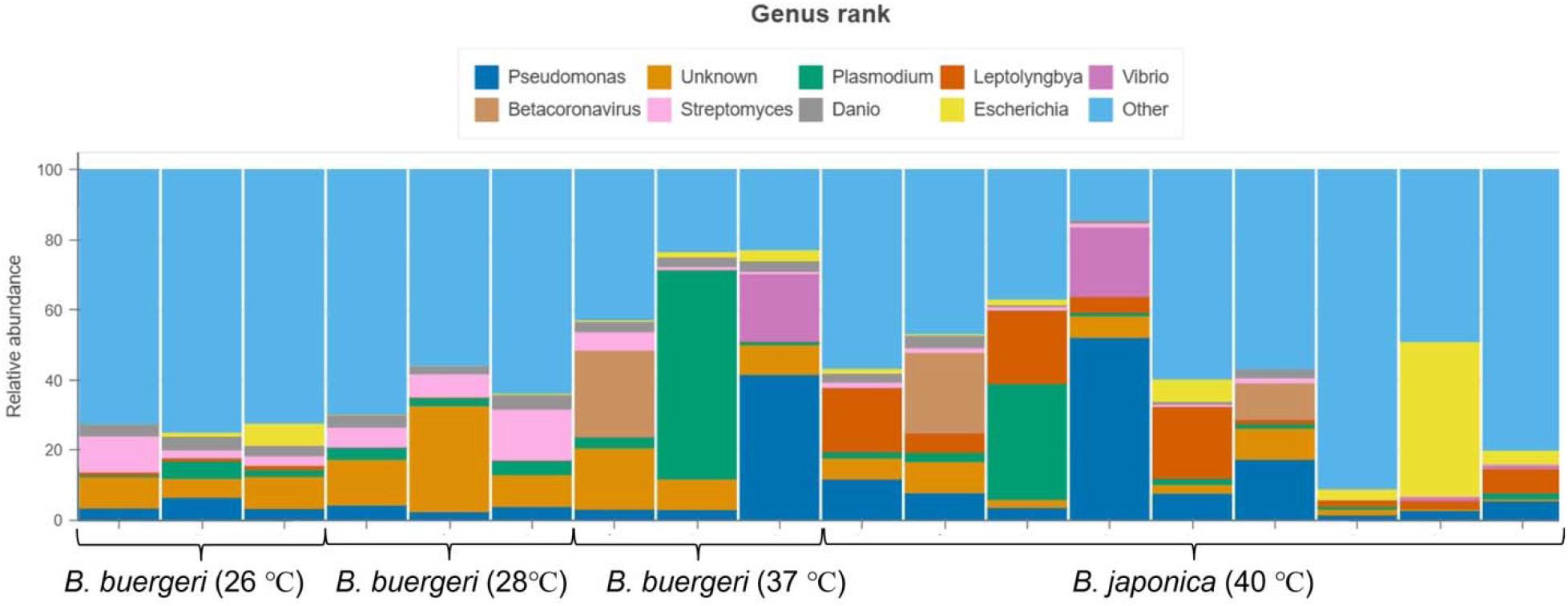
Read abundance of *B. buergeri* and *B. japonica* estimated from the Kraken2/Bracken pipeline. Barplot of the nine most abundant taxa at the genus level in all samples, where each bar represents an individual sample. Any remaining taxa were collapsed under the “Others” category to facilitate visualization. The y-axis indicates the relative abundance of each taxon as a percentage for each sample. Tables S15-S19 show the statistically significant taxa in each test.

## Discussion

### Evolutionary relationships of two novel strains of genus *Leptolyngbya*

We isolated two novel strains of *Leptolyngbya* from “Kawara-no-yukko” and “Seranma,” both of which had filamentous, nonheterocystous structures and hot spring habitats. Whole-genome sequencing revealed that the draft genomes of both strains have a single circular chromosome of approximately 6 Mb and several circular and linear contigs that are shorter than the chromosome.

Synteny plots showed that *L. boryana* had few homologous genomic regions across a contig with a length of approximately 280 kb in the draft genome of *L*. sp. Akita, suggesting that these genomic regions might be newly acquired regions in *L*. sp. Akita or a closely related strain of *Leptolyngbya*. Although there are few similar reports for other strains of *Leptolyngbya*, some strains of *Leptolyngbya* may also have genomic regions in addition to their chromosomal regions. This was consistent with the fact that a single contig of *L*. sp. Seranma and a single scaffold of *L*. sp. JSC-1 were almost homologous, in addition to their chromosomes.

Phylogenetic trees of the 16S rRNA and Unicore gene sets showed that *L*. sp. Akita and *L*. sp. Seranma was closely related to the *L. boryana* strain group and *L*. sp. JSC-1, and these trends were also observed in the BLAST-based gene composition analysis. In particular, gene composition analysis revealed that the GO terms “nitrogen compound metabolic process” and “transition metal ion binding” were enriched in the gene cluster, which was specific to the strain group of *L*. sp. Akita. All genes of *L*. sp. Akita, which had these GO terms in this gene clusters, were “NHLP leader peptide family RiPP (ribosomally synthesized and posttranslationally modified peptides) precursor”. Interestingly, these genes formed a cluster in the *L*. sp. Akita, and similar gene clusters were found in the genomes of *L. boryana* and *L*. sp. GGD. This suggests that the gene cluster of the “NHLP leader peptide family RiPP precursor,” similar to the biosynthetic gene cluster (BGC) in cyanobacteria (Yi *et al*. 2024), is a characteristic genetic element that could explain the strain group of *L*. sp. Akita. Gene composition analysis also revealed that the GO terms “DNA transposition” and “transposase activity” were enriched in the gene cluster, which was specific to the strain group of *L*. sp. Seranma. Since these genes were not significantly accumulated in *L*. sp. JSC-1, it is considered that transposon activity specifically increased in the lineage leading to *L*. sp. Seranma after divergence from *L*. sp. JSC-1. As mentioned in Yao et al. (2021) (Yao *et al*. 2021), transposons and other repetitive sequences mediate genomic rearrangements and are therefore likely involved in adaptation to diverse environments through the modification of gene expression. Thus, it is conceivable that transposon-mediated genomic structural modifications may have contributed to adaptation to high-temperature environments in *L*. sp. Seranma as well.

In contrast, no GO terms were enriched in the strain-group-specific gene clusters of *L*. sp. JSC-1, “phycobilisome” (PBS) or “photosynthesis”-related GO terms were enriched in the gene clusters, which tended to be BH (not reciprocal but best hit (i.e., simply homologous)) in the other strains, except for *L*. sp. Seranma and *L*. sp. JSC-1. Among these genes, C-phycoerythrin subunit alpha (*cpeA*) and C-phycoerythrin subunit beta (*cpeB*) were unique to the strain group containing *L*. sp Seranma, *L*. sp. JSC-1, *L*. ectocarpi, *L*. sp. Heron Island J and *L*. sp. ‘hensonii’, and the BH genes in the other strains were phycocyanin subunit alpha or phycocyanin subunit beta. A mechanism by which a part of the C-phycoerythrin family proteins composing PBS were replaced with *cpeA* and *cpeB* under far-red light has been reported in the chromatic acclimation of *L*. sp. JSC-1 (Gan *et al*. 2014). In addition to *cpeA* and *cpeB*, allophycocyanin subunit alpha-B (*apcD*), allophycocyanin subunit beta (*apcB*), photosystem I core protein (*psaA*), and photosystem I core protein (*psaB*) were also included in the strain-specific gene cluster. The different sequences and structures of these genes from homologs in other strains, with no reports of chromatic acclimation, may play an important role in chromatic acclimation. In general, these results were consistent with the phylogenetic relationships of the newly isolated strains inferred by 16S rRNA and Unicore, and also indicated that our gene composition analysis could identify characteristic genes in the targeted strain group.

Consistent with the comparative genomic analysis, the absorption spectral responses revealed a clear difference in light-acclimation capacity between *L*. sp. Akita and *L*. sp. Seranma. While *L*. sp. Akita exhibited largely invariant absorption spectra under different light conditions, indicating a relatively fixed photosynthetic architecture, *L*. sp. Seranma showed pronounced, light-dependent changes in absorption profiles, reflecting substantial spectral plasticity and adaptive reorganization of its light-harvesting system. Although phycobilisome-related genes were conserved among some *Leptolyngbya* strains, the *rfpABC* gene cluster encoding the master regulators of far-red light photoacclimation was detected only in *L*. sp. Seranma and *L*. sp. JSC-1 (data not shown). This distribution is consistent with the absence of chromatic acclimation in *L*. sp. Akita and with the lack of previous reports of chromatic acclimation in other *Leptolyngbya* strains. Together, these results suggest that the expression of chromatic acclimation requires not only the presence of phycobilisome-associated genes but also the conservation of regulatory modules such as the *rfp* cluster within the genome.

### Potential roles of lineage-specific genes in thermal stress adaptation of *L*. sp. Akita and the *L*. sp. Seranma–*L*. sp. JSC-1 lineage

The present analysis identified a set of gene annotations that were conserved among hot-spring-derived strains but absent from non-thermal environment-derived strains, suggesting that these genes may represent lineage-specific acquisitions associated with adaptation to high-temperature environments. Although direct functional validation is lacking, the functional categories enriched among these genes provide insight into the potential mechanisms underlying thermal stress adaptation in these lineages.

Among the genes shared between *L*. sp. Akita and *L*. sp. Seranma, several were annotated as oxidative stress response factors, including peroxiredoxin family proteins (Fiorentino *et al*. 2020; Bolduc *et al*. 2021) and Dyp-type peroxidases (Ogola *et al*. 2009). Elevated temperatures are known to exacerbate the production of reactive oxygen species, even in photosynthetic or aerobic microorganisms, leading to oxidative damage to proteins, lipids, and nucleic acids. These oxidative stress–related genes suggests that reinforced detoxification of reactive oxygen species may represent a common adaptive strategy in hot-spring-derived *Leptolyngbya* strains.

In addition to oxidative stress–related genes, the shared gene set also included membrane stress response–related factors, such as M23 family metallopeptidases (Razew *et al*. 2022), NfeD family proteins (Ukleja *et al*. 2024), and dynamin family proteins (Sattler and Graumann 2022). High temperatures can compromise membrane integrity and protein localization, particularly in organisms inhabiting thermally fluctuating environments. These genes may therefore contribute to maintaining membrane stability or facilitating membrane remodeling under heat stress, consistent with observations in other thermophilic and thermotolerant microorganisms (Yamada 2024).

Notably, the shared gene set further included RNA remodeling factors such as DEAD/DEAH box helicases. Temperature changes profoundly affect RNA secondary structure, which in turn influences translation efficiency and RNA stability. DEAD/DEAH box helicases are known to function as ATP-dependent RNA remodeling enzymes that maintain RNA functionality under stress conditions. Although many DEAD-box RNA helicases have been primarily characterized as cold shock–responsive factors, previous studies have shown that the loss of specific DEAD/DEAH box RNA helicases can also reduce the maximum permissive growth temperature in *Escherichia coli* (Markkula *et al*. 2012), indicating that individual members of this family may contribute to growth across a broader temperature range. These putatively acquired genes suggests that post-transcriptional regulation and RNA quality control may play an important role in thermal adaptation in these lineages. Considering the similarity of their habitats, particularly exposure to high-temperature environments, these novel members of shared gene families suggest that functionally analogous stress response factors were independently acquired or diversified during lineage divergence. Such a process likely reflects convergent evolutionary trajectories associated with adaptation to thermal stress, rather than simple inheritance from a common ancestor.

Genes detected exclusively in *L*. sp. Akita were included several factors potentially related to DNA protection and genome maintenance, including FtsK/SpoIIIE domain-containing proteins (Begg, Dewar and Donachie 1995), RusA family crossover junction endodeoxyribonucleases (Mahdi *et al*. 1996), and UvrD-like helicases (Epshtein *et al*. 2014). Elevated temperatures can increase DNA damage and replication stress, and the acquisition of such genes may enhance the capacity of *L*. sp. Akita to deal with DNA stress in their thermal environments. In addition, the presence of AAA family ATPases and AAA domain-containing proteins (Weibezahn *et al*. 2004; Kamata *et al*. 2005; Zolkiewski, Zhang and Nagy 2012) suggests a potential reinforcement of protein quality control systems, including ATP-dependent disaggregation or proteolytic pathways, which are critical for mitigating heat-induced protein damage.

In contrast, genes uniquely detected in *L*. sp. Seranma included oxidative stress response enzymes, protein repair enzymes such as protein-L-isoaspartate O-methyltransferase (Kindrachukt *et al*. 2003), and chaperone regulatory factors such as CbpM (Chenoweth, Trun and Wickner 2007). Protein-L-isoaspartate O-methyltransferase functions in the repair of damaged proteins that accumulate under prolonged stress conditions, while CbpM modulates the activity of the DnaK–DnaJ chaperone system. Together, these genes suggest that *L*. sp. Seranma may rely more strongly on protein repair and fine-tuned regulation of chaperone activity, rather than solely on degradation-based quality control mechanisms, to maintain proteostasis under thermal stress.

Taken together, the lineage-specific distribution of these genes indicates that thermal stress adaptation in hot-spring-derived *Leptolyngbya* strains is likely achieved through a combination of oxidative stress mitigation, membrane stabilization, RNA remodeling, and protein quality control, with distinct lineages emphasizing different components of this multifaceted stress response. While these genes cannot be regarded as direct determinants of thermotolerance, their consistent association with hot-spring-derived lineages suggests that they may collectively enhance cellular robustness in high-temperature environments.

Future experimental validation, including expression analyses under heat stress and functional characterization of selected candidate genes, will be required to clarify the precise contributions of these lineage-specific genes to thermal adaptation.

### Ecological relationships of cyanobacteria and sympatric animals and utilization as functional food resources

Finally, we focused on the high-temperature tolerance of *B. buergeri* and *B. japonica* tadpoles that sympatrically lived in their habitats, hot-springs of “Kawahara-no-Yukko” and “Seranma”, and compared their food habits using intestinal metagenomic analysis. The results of the GLM tests suggested that the genera *Letolyngbya* and *Limnothrix* were the characteristic parameters that could explain the difference between the food habitats of *B. buergeri* and *B. japonica* tadpoles. Similarly, Wilcoxon tests supported the genus *Leptolyngbya* as a parameter that could indicate differences in food habitats between *B. buergeri* and *B. japonica* tadpoles. Considering that reads of the genus *Letolyngbya* tend to be more frequently detected in *B. japonica* tadpoles and that a few reads of the genus *Limnothrix* were only detected in *B. buergeri* tadpoles derived from 27 °C, it was suggested that *Leptolyngbya* could more strongly explain the difference between *B. buergeri* and *B. japonica*, especially more frequently used food resources. Unfortunately, in the other genera that showed statistical significance between *B. buergeri* tadpoles derived from each temperature, the number of reads was smaller than that of the genus *Leptolyngbya*, and the differences in the food habits of *B. buergeri* under each temperature environment were unclear. In addition, it is important to note that parametric hypothesis tests detected no statistical significance, whereas nonparametric hypothesis tests did. These results suggested that the difference could not be detected in the food habitats of tadpoles if we assumed that the read abundance of the genus *Leptolyngbya* followed a normal distribution. In other words, each tadpole might use *Leptolyngbya* as a temporary food source rather than a regular food source. *B. buergeri* and *B. japonica* tadpoles collected from these hot-springs did not show clear high-temperature tolerance in the rearing experiments, so the high-temperature tolerance of these tadpoles might have high phenotypic plasticity (data not shown), and these temporary food resources might cause the plasticity, especially in *B. japonica*. Overall, bacterial mats containing *Leptolyngbya* might allow amphibians that prefer low-temperature environments to obtain the new adaptive trait of high-temperature tolerance; however, it was difficult to prove that the amount of *Leptolyngbya* as their food directly gave them high-temperature tolerance. Thus, we plan to perform feeding experiments on tadpoles using isolated and cultured *Leptolyngbya* to verify these mechanisms in the future.

## Acknowledgement

We would like to express our sincere gratitude to Dr. Makoto Suzuki, Amphibian Research Center, Hiroshima University, for his assistance with tadpole sampling in Akita, Japan. Additionally, we extend our appreciation to Dr. Takuya Aoyanagi and Dr. Shin-ichi Aoki, Institute for Microalgal Technology, for their cooperation in culturing the isolated cyanobacteria, which greatly contributed to the success of this research.

