## Supplemental Figures for "Comparative genomic analysis of cyanobacteria as amphibian food sources: insights into high-temperature tolerance potential"

**Supplementary informations**

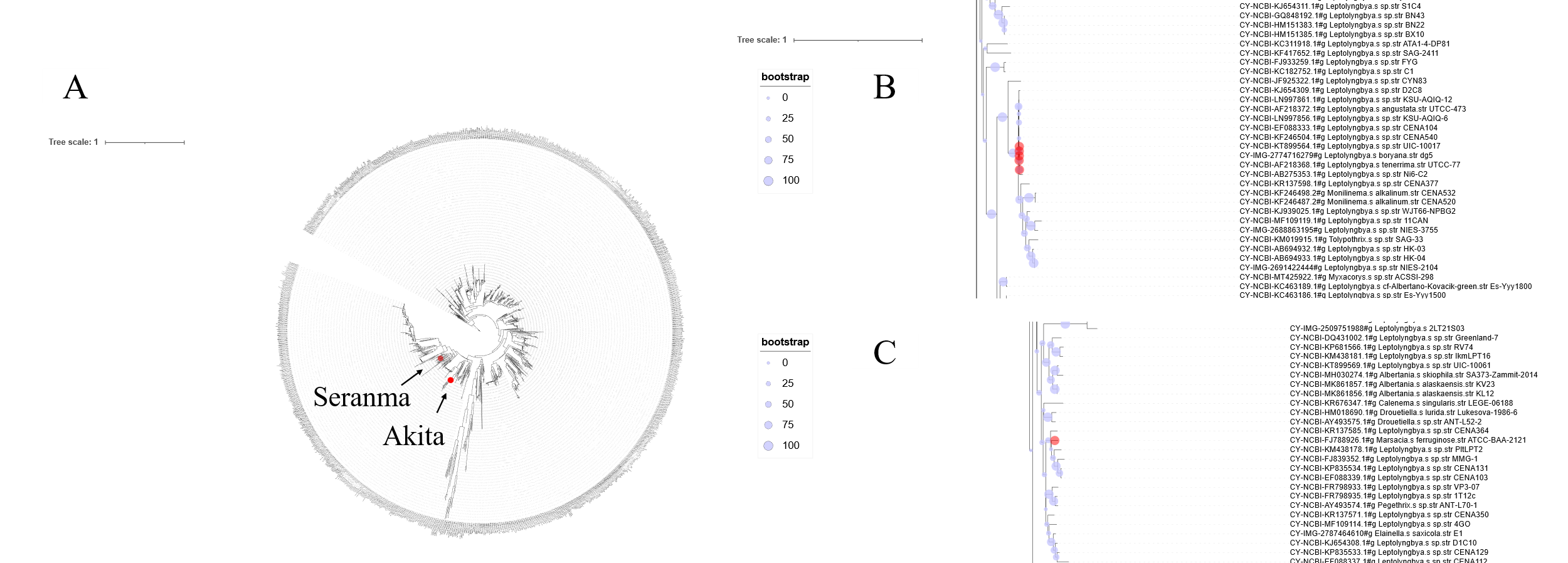

Fig. S1 A phylogenetic tree generated from Cydrasil showed (A) the location of *L*. sp. Akita and *L*. sp. Seranma in the whole tree, (B) branch of *L*. sp. Akita, and (C) branch of *L*. sp. Seranma

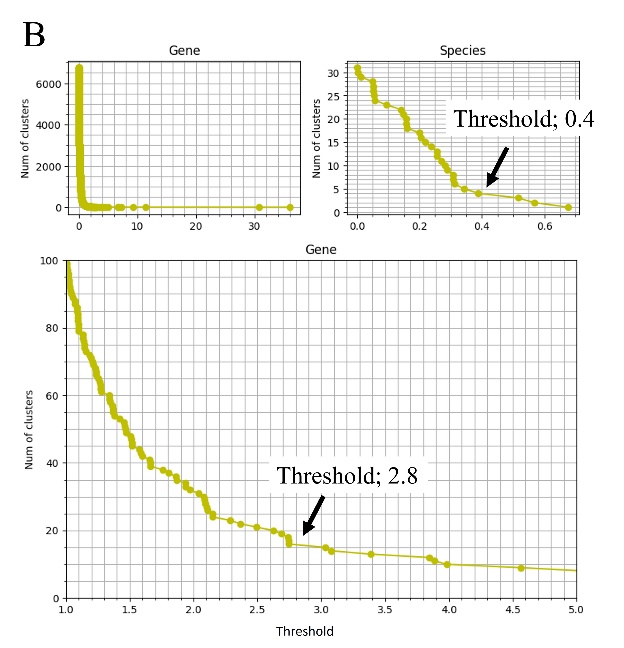

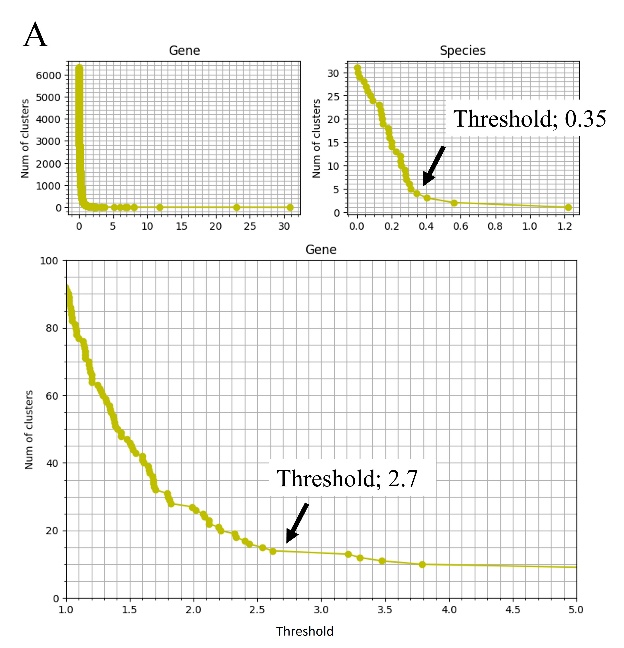

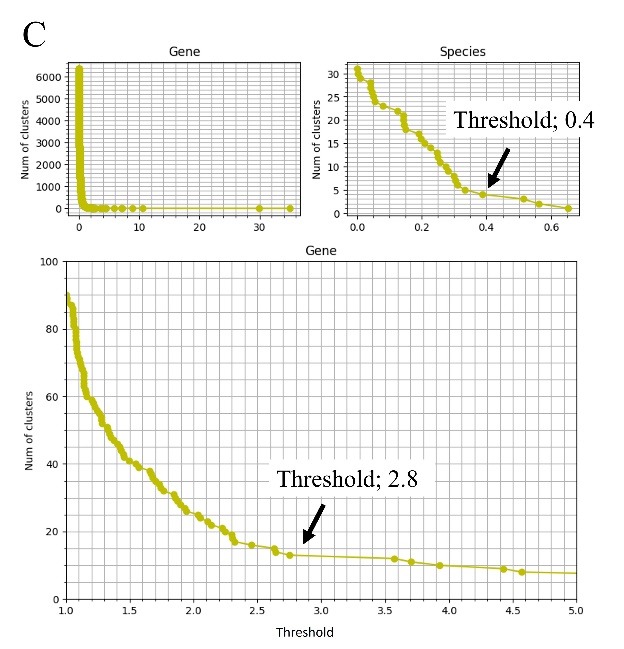

Fig. S2

Thresholds of hierarchical clustering in (A) *L*. sp. Akita, (B) *L*. sp. Seranma, and (C) *L*. sp. JSC-1

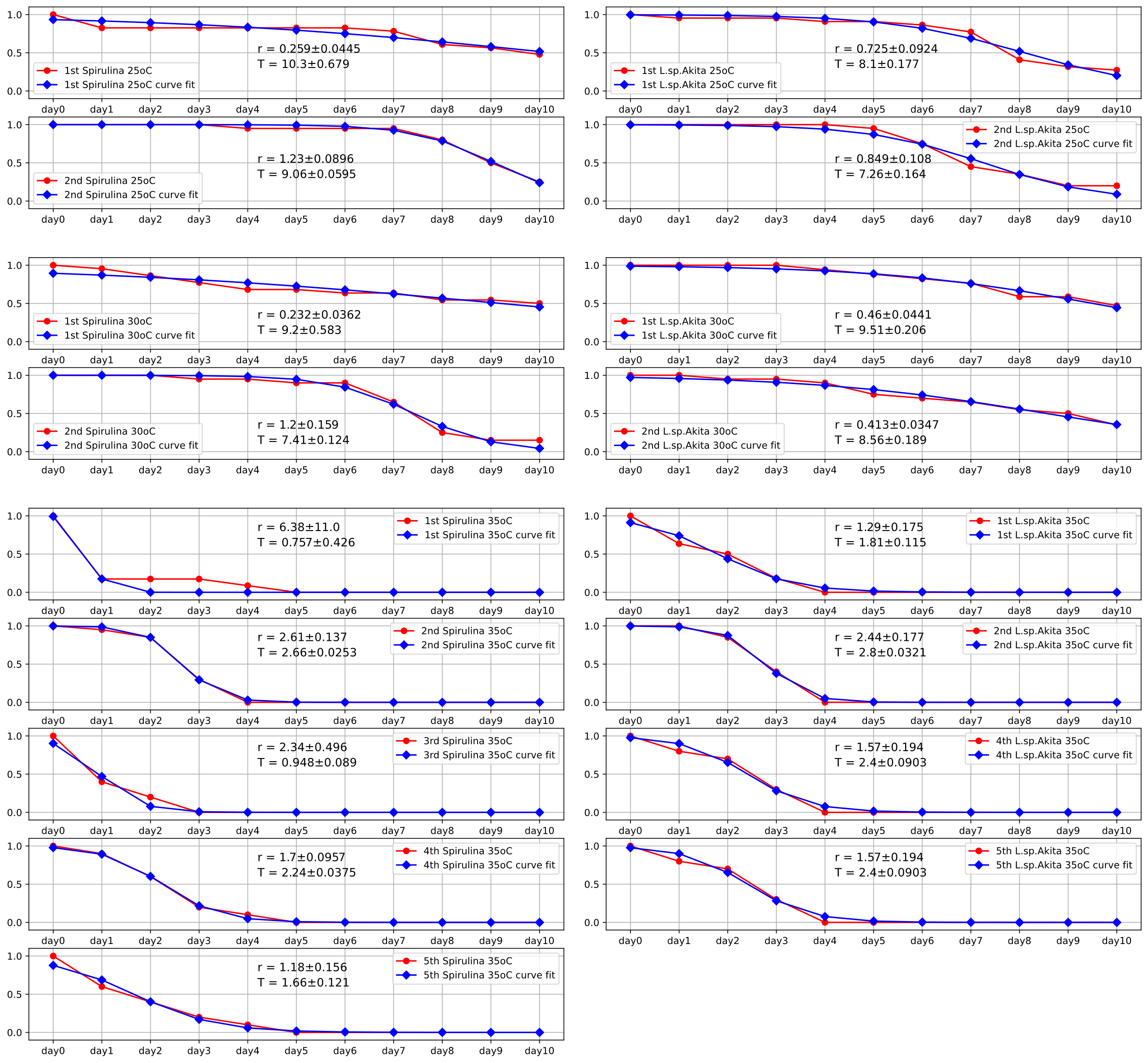
Fig. S3

Time series of survival rates (red) and the sigmoid functions with the estimated parameters (blue) and their standard deviations in all trials.

Table S1 Metrics in draft genome of each *Leptolyngbya* strain used in this study

Table S2 Best hit protein IDs and categories of reciprocal BLASTP search between predicted proteins of *L.* sp. Akita and those of each *Leptolyngbya* strain

Table S3 Best hit protein IDs and categories of reciprocal BLASTP search between predicted proteins of *L.* sp. Seranma and those of each *Leptolyngbya* strain

Table S4 Best hit protein IDs and categories of reciprocal BLASTP search between predicted proteins of *L.* sp. JSC-1 and those of each *Leptolyngbya* strain

Table S5 Cluster IDs of each protein in *L*. sp. Akita, *L*. sp. Seranma, and *L*. sp. JSC-1

Table S6 Results of InterProScan (Pfam) in *L*. sp. Akita

Table S7 Results of InterProScan (Pfam) in *L*. sp. Seranma

Table S8 Results of InterProScan (Pfam) in *L*. sp. JSC-1

Table S9 Significantly enriched GO terms in each gene cluster on *L.* sp. Akita

Table S10 Significantly enriched GO terms in each gene cluster on *L.* sp. Seranma

Table S11 Significantly enriched GO terms in each gene cluster on *L.* sp. JSC-1

Table S12 Results of Kruscal-Wallis test and glm test between read abundances of *B. buergeri* of 26 °C, *B. buergeri* of 28 °C, *B. buergeri* of 37 °C, and *B. japonica* of 40 °C

Table S13 Results of Welch's t-test and Wilcoxon test between read abundances of *B*. *buergeri* and *B*. *japonica*

Table S14 Results of Welch's t-test and Wilcoxon test between read abundances of *B. buergeri* of 26 °C and *B. buergeri* of 28 °C

Table S15 Results of Welch's t-test and Wilcoxon test between read abundances of *B. buergeri* of 26 °C and *B. buergeri* of 37 °C

Table S16 Results of Welch's t-test and Wilcoxon test between read abundances of *B. buergeri* of 28 °C and *B. buergeri* of 37 °C

Table S17 Survival rates of *Xenopus tropicalis* tadpoles in all trials of feeding experiments.

TableS18 Results of Welch's t-test between the average survival rates in each time point

Table S19 Estimated mortality rates (r) and inflection points (T) of the sigmoid functions in all trials.

Table S20 Results of Welch's t-test between estimated parameters in each combination of trials.
